# Whole-plant optimality predicts changes in leaf nitrogen under variable CO_2_ and nutrient availability

**DOI:** 10.1101/785329

**Authors:** Silvia Caldararu, Tea Thum, Lin Yu, Sönke Zaehle

**Author notes:** Corresponding author: Silvia Caldararu.

## Abstract

- Vegetation nutrient limitation is essential for understanding ecosystem responses to global change. In particular, leaf nitrogen (N) is known to be plastic under changed nutrient limitation. However, models can often not capture these observed changes, leading to erroneous predictions of whole-ecosystem stocks and fluxes.
- We hypothesise that an optimality approach can improve representation of leaf N content compared to existing empirical approaches. Unlike previous optimality-based approaches, which adjust foliar N concentrations based on canopy carbon export, we use a maximisation criteria based on whole-plant growth and allow for a lagged response of foliar N to this maximisation criterion to account for the limited plasticity of this plant trait. We test these model variants at a range of Free-Air CO_2_ Enrichment (FACE) and N fertilisation experimental sites.
- We show a model solely based on canopy carbon export fails to reproduce observed patterns and predicts decreasing leaf N content with increased N availability. However, an optimal model which maximises total plant growth can correctly reproduce the observed patterns.
- The optimality model we present here is a whole-plant approach which reproduces biologically realistic changes in leaf N and can thereby improve ecosystem-level predictions under transient conditions.

## 1 Introduction

The response of plants to changes in environmental conditions such as climate, atmospheric CO_2_ concentration or nutrient inputs is dependent on the plants’ ability to acclimate and adapt to the new conditions. In particular, there is strong evidence that plants can respond plastically to changes in nutrient limitation (Bloom *et al*., 1985; Chapin *et al*., 1986), which in turn affects their response to elevated atmospheric CO_2_ (Ainsworth & Long, 2005; Long *et al*., 2004). However, such processes are lacking or poorly represented in terrestrial biosphere models (TBMs) (Prentice *et al*., 2015; Medlyn *et al*., 2015; Zaehle *et al*., 2014). Therefore, in this study we set out to provide a better representation of plant, and specifically canopy, response to increasing nutrient limitation.

One of the main roles of nitrogen (N) in plant physiology is as a component of proteins and specifically enzymes that form part of the photosynthetic apparatus (Hawkesford *et al*., 2012; Evans, 1989), which is why leaves create the highest growth demand for N. Therefore, understanding leaf N physiology and its response to changing environmental drivers is essential for understanding whole-plant and ecosystem level response to N limitation.

A decrease in leaf N content has been observed over the last decades and has been previously attributed to increasing atmospheric CO_2_ (Jonard *et al*., 2015; McLauchlan *et al*., 2010). A similar decrease has also been observed under experimentally elevated CO_2_ (Ellsworth *et al*., 2004). On the other hand, increased leaf N concentration occurs in nitrogen addition experiments (e.g. Magill *et al*., 2004; Houle & Moore, 2008; McNulty *et al*., 2005; Sikström, 2002) and under high N deposition (Fleischer *et al*., 2013; McNeil *et al*., 2007). There have been a number of explanations proposed for this observed plasticity, including both plant N use and changes in the photosynthetic apparatus. Firstly, plants increase their leaf N under increased N availability to make better use of available resources. Under elevated CO_2_, plants can achieve the same carbon assimilation rate with a lower leaf N content (Stitt & Krapp, 1999; Drake *et al*., 1997). Additionally, there are other associated changes in the photosynthetic apparatus under elevated CO_2_ which can contribute to the observed changes in leaf N (Ainsworth & Long, 2005; Rogers & Humphries, 2000). Another proposed mechanism for the observed decline in leaf N content under elevated CO_2_ is known as the dilution effect, an accumulation of non-structural carbohydrates in the leaves which cause an overall lower N concentration, although there is little experimental evidence for this process playing an important role (Feng *et al*., 2015; Wujeska-Klause *et al*., 2019). Furthermore, the extent to which the leaf N content changes depends on the initial N limitation status of the plant (Ainsworth & Long, 2005; Stitt & Krapp, 1999).

From a modelling perspective, TBMs which include N processes represent the leaf N content either as a fixed parameter or as a flexible value which responds to plant N limitation, generally following an empirical function (Zaehle & Dalmonech, 2011; Thomas *et al*., 2013a,b). Meyerholt & Zaehle (2015) show that models which include a flexible C:N ratio have better predictive capability than models with a fixed ratio when compared to observations from N fertilisation experiments. However, they also found that models with flexible stoichiometry tend to overestimate the increase in foliar N under N fertilisation, thereby increasing the N costs of new tissue, leading to a lower than observed growth response. A multi-model comparison at two FACE sites has shown that models with flexible stoichiometry can overestimate the leaf N response to elevated CO_2_, in some cases reaching unrealisticly low values (Zaehle *et al*., 2014). These two results together point to a need for a better representation of dynamic leaf N to capture the N dynamics of ecosystems under changing environmental conditions.

Optimality principles have long been proposed as a theoretical way of representing plant plasticity (Givnish, 1988; Mäkelä *et al*., 2002). The underlying hypothesis is that plants alter their physiology or morphology to maximise growth over their lifetime. The concept has been previously applied to individual processes such as photosynthesis (as the coordination hypothesis Medlyn, 1996; Maire *et al*., 2012; Ali *et al*., 2015; Smith *et al*., 2019), stomatal conductance (Cowan & Farquhar, 1977; Medlyn *et al*., 2011), biomass allocation (Mäkelä *et al*., 2008), nutrient content (Dewar, 1996) and phenology (Caldararu *et al*., 2014). Optimality has the potential to improve the representation of vegetation processes in TBMs by providing a way to include complex responses that can accurately capture observations without a large number of additional parameters. A number of studies have built basic whole-plant models centred around the optimality hypothesis (Franklin, 2007; McMurtrie *et al*., 2008). More recently, optimality concepts, and specifically leaf-level coordination of photosynthetic parameters, have started to be incorporated into full TBMs (e.g. Ali *et al*., 2015; Haverd *et al*., 2018). However, there are still a number of open questions about the exact mode of implementation, including which processes and variables to optimise and the timescale of this optimisation (Dewar *et al*., 2009).

Here, we implement an optimality-based method for representing changes in leaf N within a newly developed TBM, QUINCY (QUantifying Interactions between terrestrial Nutrient CYcles and the climate system, Thum *et al*., 2019). A TBM, with its detailed representation of soil and vegetation processes, allows us to identify the drivers and assess the implications, of changes in leaf stoichiometry given the ecosystem-scale nutrient balance. In particular, the detailed representation of canopy and photosynthetic processes allows us to go beyond the basic optimality models used in the past and predict time-variant leaf N content and not simply at-equilibrium values. Instead of relying on the assumption that plant properties (here leaf N) are optimal at any timestep, we rely on a dynamic, towards equilibrium approach, in which plants adjust their properties (here leaf N) at each model timestep towards the optimal value given average environmental conditions over a predefined period. This approach reflects the known limitations to plant plasticity and thereby provides a solution to the timescale of optimisation problem introduced above. On the other hand, the use of optimality theory allows us to leverage basic processes already existent within the TBM and therefore represent complex plastic behaviour without introducing a large number of parameters in addition to those already present in the model.

In this study, we test the hypothesis that changes in leaf N concentrations can be explained by two main drivers: (1) the limitation to growth by N availability caused by an increase in leaf N content and, (2) the increase in carbon export, and thus growth, through an increase in leaf N content. The first driver is the one used by existing models to describe variations in leaf N, while the second is commonly used in optimality approaches. To explore these two drivers we use four different model setups for representing leaf N: fixed leaf N content, empirical (which includes only the N availability criteria), optimal C export (which includes only the maximum C export criteria) and optimal growth (which includes both). We test the effect of these different formulations on plant growth and foliar N responses to changes in CO_2_ and N availability using the resulting model at two FACE sites and a range of N fertilisation sites.

## 2 Methods

### 2.1 Model description

#### 2.1.1 The QUINCY terrestrial biosphere model

We implement the above options in a terrestrial biosphere model, QUINCY, described in detail in Thum *et al*. (2019). Here we briefly present the processes related to the representation of dynamic leaf N, followed by a detailed description of the optimality implementation.

QUINCY represents fully coupled C, N and P as well as water and energy cycles (Fig. 1). The model employs a multi-layer canopy scheme, which includes a representation of photosynthesis and canopy conductance, for sunlit and shaded leaves separately within every layer. Total canopy N is distributed to each layer, with exponentially decreasing N content towards the bottom of the canopy. Variations in average leaf N content (described below) are propagated to each canopy layer. Photosynthesis is represented following the model of Kull & Kruijt (1998). Leaf N is separated into structural and photosynthetic (chlorophyll, Rubisco, and electron transport) fractions, each of which is directly related to parameters in our photosynthesis model (Kull & Kruijt, 1998; Zaehle & Friend, 2010). The structural N fraction is expressed as a linear function of leaf N content, decreasing with higher leaf N (Evans, 1989; Friend *et al*., 1997). The fraction of photosynthetic N allocated to chlorophyll increases with canopy depth, while the Rubisco and electron transport components are adjusted to give a constant ratio of the respective photosynthetic parameters (*V_cmax,_*_25_ and *J_max,_*_25_. Maintenance respiration in the model is represented as a linear function of specific tissue N. Growth respiration is calculated as being proportional to the amount of new tissue built. Both photosynthesis and maintenance respiration acclimate to growth temperature following Friend (2010) and Atkin *et al*. (2014).

**Figure 1:**
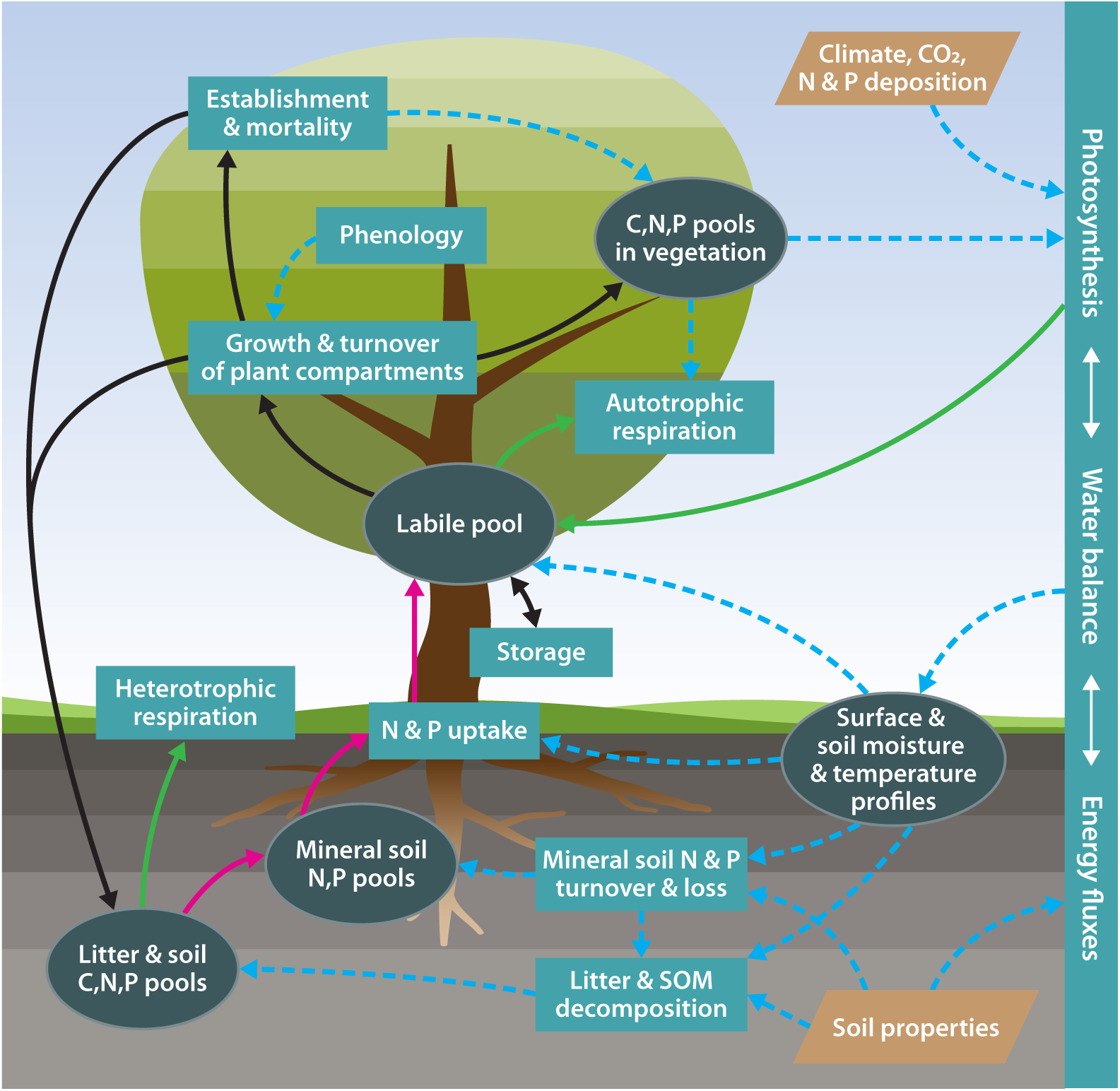
Schematic representation of the QUINCY baseline model structure. Elipses: biogeochemical pools and other state variables; rectangles: biogeochemical processes; tetraeders: model input; solid green lines: carbon fluxes; solid dark red lines: nitrogen and phosphorus fluxes; solid black lines: carbon, nitrogen and phosphorus fluxes; dotted blue lines: effects (Thum *et al*., 2019).

Nutrient uptake is calculated as a function of fine root biomass, soil inorganic nutrient content and plant demand for each specific nutrient, where the demand depends on the ratio of available to required nutrients for growth.

Photosynthesis and growth processes are separated in QUINCY by the introduction of two non-structural pools, the labile and reserve pool. Carbon assimilated through photosynthesis and nutrients taken up by the roots enter the labile pool, and are subsequently allocated to new tissue growth, transferred to the reserve compartment, or in the case of carbon, used for respiration. Growth follows a functional balance approach based on an extended pipe-theory representation, generating size-dependent allometric relationships between leaf, fine root, coarse root, and sapwood mass. Actual growth is constrained by available nutrients and tissue-specific stoichiometry. Stand-level vegetation dynamics is represented as density-dependent mortality and establishment based on the existing seedbed, as the model explicitly allocates to a reproductive pool. All pools and fluxes are representative of an average individual.

The QUINCY model includes a detailed representation of soil carbon and nutrient pools, the details of which can be found in Thum *et al*. (2019).

#### 2.1.2 Tissue stoichiometry

The N content of leaves and fine roots at each half-hourly timestep Δ*t* is updated given a direction variable, *D_N_* (unitless), and a parameter representing the maximum rate of change, *δ_N_* (day*^−^*^1^), so that:

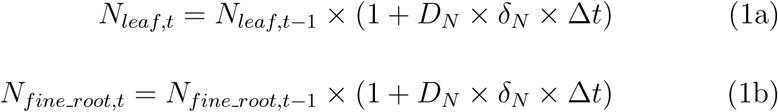

The *N_leaf_* values here refer to canopy average values of leaf N content. To conserve the mass balance, new tissue is added with a leaf N equal to that calculated above, while old tissue changes its N content towards the target C:N by recycling N through the labile pool at a timescale of 10 days. All values of leaf N content below refer to this target average leaf N value.

The C:N ratio of fine roots is represented as being directly proportional to that of leaves (Zaehle & Friend, 2010). We consider the N content of the woody and fruit pools to be constant (Meyerholt & Zaehle, 2015). Even though the model variants described below are centred on changes in leaf N, the associated changes in fine root N need to be taken into account at the whole plant level and will be calculated throughout. Note that as in our model the specific leaf area (SLA) remains constant, a fractional change in total leaf N results in an equal fractional change in C:N ratio and leaf N content per mass and per area.

Below, we describe in detail the four model options for calculating foliar stoichiometry, which in essence are four options for calculating the value of *D_N_*. The rate of change parameter, *δ_N_* is the same for all variants.

#### 2.1.3 Variant 0: fixed leaf N

For the basic model variant, tissue stoichiometry is kept constant and the value of *D_N_* is simply set to zero. Stoichiometry parameters are prescribed, following PFT-specific values from the TRY trait data base (Kattge *et al*., 2011).

#### 2.1.4 Variant 1: empirical leaf N

The empirical variation is based on Zaehle & Friend (2010). The leaf N concentration is modified based on the relative availabilities of labile carbon (*C_labile_*) and nitrogen (*N_labile_*) for growth, given the stoichiometric requirement, i.e. the C:N ratio *χ^CN^_growth_*. The direction of change variable, *D_N_*, is calculated as:

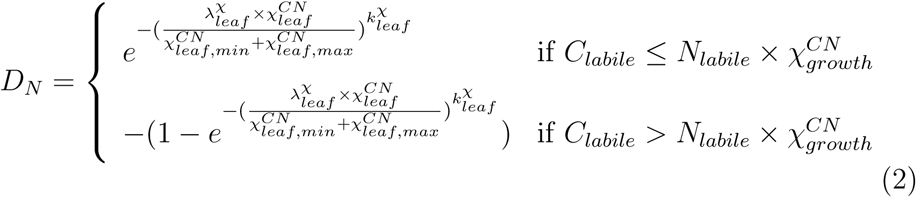

In the above, *λ^χ^_leaf_* and *k^χ^_leaf_* are empirical shape parameters, *χ^CN^_leaf_* is the current leaf stoichiometric ratio and *χ^CN^_leaf,min_* and *χ^CN^_leaf,max_* are PFT-specific parameters, taken from the TRY database (Kattge *et al*., 2011). The C:N ratio of new growth *χ^CN^_growth_* is a variable calculated at every timestep given fractions allocated to each pool and current stoichiometry of all pools. Note that unlike the two optimality schemes described below, here *D_N_* can take values between -1 and 1 but the same notation has been kept for consistency.

#### 2.1.5 Variant 2: optimal C export

Our first model assumption is that plants modify their leaf N to maximise net C assimilation. Increased leaf N will increase photosynthesis (gross assimilation, *A_g_*) but also increase tissue maintenance respiration, resulting in a peaked relationship between net assimilation (*A_n_*) and leaf nitrogen, as shown in Fig. 2. Due to the non-linear nature of N distribution in the canopy and the photosynthesis model used in QUINCY, we calculate the optimal leaf N for which net carbon export is maximised through a numerical approach, as follows.

**Figure 2:**
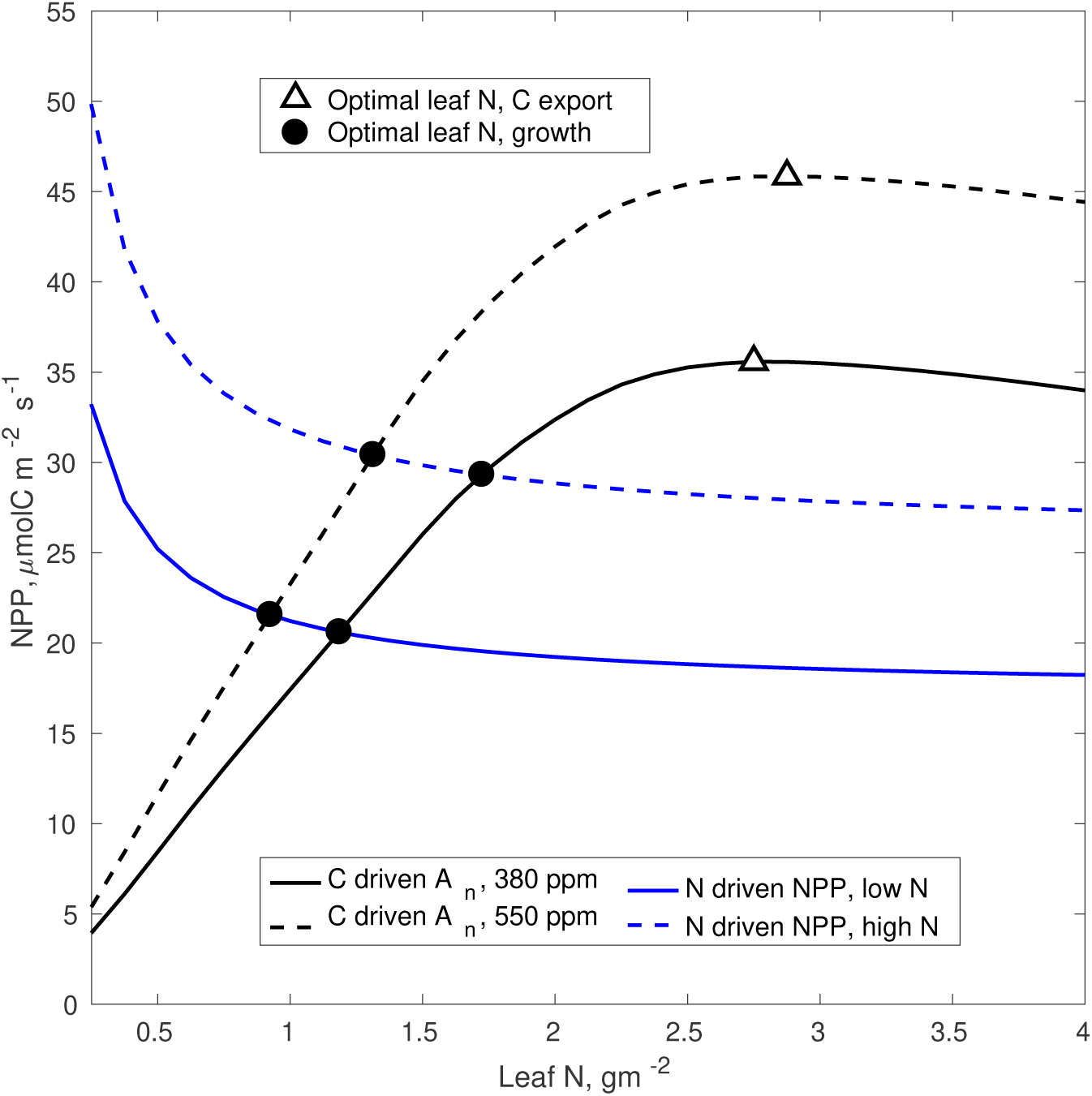
Theoretical representation of optimal leaf N. Black lines show the variation in net C assimilation with leaf N and blue lines show the C-equivalent plant growth given an amount of N available in the soil. Both *A_n_*and *NPP* represent whole-canopy values. *A_n_*, net assimilation refers to gross assimilation minus canopy maintenance respiration, while *NPP* refers to whole-plant net primary productivity available for growth. Leaf N values are canopy averages. Triangles show the leaf N for which maximum NPP occurs, equivalent to the optimal C export variant. Black circles show leaf N for which C uptake is equal to N uptake, equivalent to the optimal growth variant.

We calculate the direction in which the leaf N content needs to be modified by increasing the current leaf N content *N_leaf,t−_*_1_ by a given amount *δ_N_* and the fine root N content by the equivalent amount to obtain two new values, *N_leaf,δ_* and *N_fine_root,δ_*.

We then calculate the net assimilation *A_n,δ_* given the new *N_leaf,δ_*as:

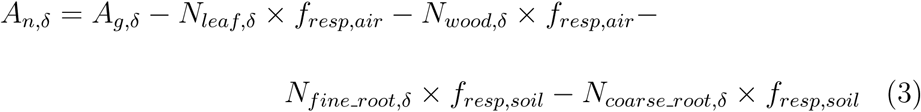

where *f_resp,air_* and *f_resp,soil_* are the maintenance respiration rate per unit N given the temperature of the air and soil respectively. *A_g,δ_* is the gross canopy assimilation given the new leaf N, calculated for each canopy layer. As the C:N ratio of wood and coarse roots is considered to be constant, the difference in *A_n_*, *dA_n_* resulting from the change in tissue N, *dN_leaf_* is equal to the change in photosynthesis with the change in N, minus the change in tissue maintenance respiration, *dR_leaf_* and *dR_fine_*___*_root_*, with the change in N content of each specific tissue:

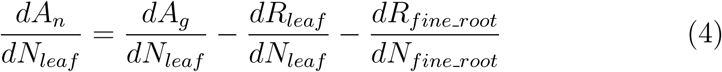

The *A_n_* values are calculated given the average meteorological conditions over a given time period, *τ_N_* (30 days). Note that this approach means that plants do not reach absolute optimality but there is a rate of change in the optimal direction, with the parameter *δ_N_* denoting the maximum amount by which the leaf N content can change in a timestep (Table 1).

**Table 1:** Model parameters introduced for each of the four variants of representing leaf N content. Parameters which are part of the baseline QUINCY model and are relevant for the processes described here can be found in Table S1. BS = broadleaf deciduous, NE = needleleaf evergreen

| Symbol | Description | Value BS | Value NE | Unit | Citation |
| --- | --- | --- | --- | --- | --- |
| **Fixed** |  |  |  |  |  |
| $\chi_{leaf}^{CN}$ | Foliar C:N | 22.5 | 39.7 | $\frac{molC}{molN}$ | Kattge *et al.* (2011) |
| **Empirical** |  |  |  |  |  |
| $\lambda_{leaf}^x$ | Shape parameter in leaf stoichiometry nutrient response | 2.0 | 2.0 | - | Zaehle & Friend (2010) |
| $k_{leaf}^x$ | Shape parameter in leaf stoichiometry nutrient response | 8.0 | 8.0 | - | Zaehle & Friend (2010) |
| $\chi_{leaf,min}^{CN}$ | Minimum foliar C:N | 14.0 | 24.0 | $\frac{molC}{molN}$ | Kattge *et al.* (2011) |
| $\chi_{leaf,max}^{C:N}$ | Maximum foliar C:N | 38.7 | 64.9 | $\frac{molC}{molN}$ | Kattge *et al.* (2011) |
| $\delta_N$ | Maximum rate of change in foliar N content | 0.0048 | 0.0048 | $day^{-1}$ | This study |
| $\tau_N$ | Response timescale for changes in foliar N content | 30 | 30 | days | This study |
| **Optimal C export & growth** |  |  |  |  |  |
| $\delta_N$ | Maximum rate of change in foliar N content | 0.0048 | 0.0048 | $day^{-1}$ | This study |
| $\tau_N$ | Response timescale for changes in foliar N content | 30 | 30 | days | This study |

We then calculate the direction variable *D_N_* so that the new actual leaf N increases for a positive return in *A_n_* and decreases for a negative return:

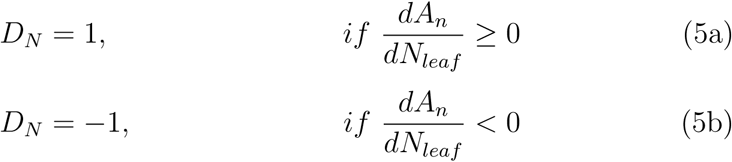

The optimal direction resulting from this criterion will vary with environmental conditions (e.g. temperature, water availability), atmospheric CO_2_ concentration (see Fig. 2), as well as light environment in the canopy as given by direct and diffuse light levels at the top of the canopy but also the amount of leaf area (Fig. S3).

#### 2.1.6 Variant 3: optimal growth

In addition to the optimal C export criterion, here we introduce an additional condition for calculating the *D_N_* direction variable, based on the relationship between the potential N-limited growth, *N_growth_*, and the potential C-limited growth, *C_growth_*. The C-limited growth can be calculated as:

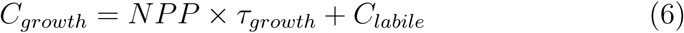

where *τ_growth_* is the timescale of plant growth, in this case equal to one year, and *C_labile_*is the C content of the labile pool, i.e. the C available for immediate growth. Note that the NPP value here is not the same as *A_n_* in previous equations as it also takes into account growth respiration and storage fluxes into the reserve pool. The N-limited growth term is calculated given the available N and the stoichiometry of new tissue, *χ^CN^_growth_*:

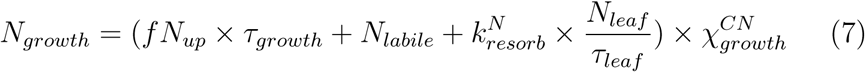

Here, *f N_up_* refers to the N uptake by the roots, *k^N^_resorb_* is a parameter denoting the rate of nutrient resorption from senescing leaves and *τ_leaf_* is the turnover time of the leaves. The term *χ^CN^_growth_*converts *N_growth_* into C units, to allow comparison with *C_growth_*. All parameter values can be found in Table S1. Both C and N availability is calculated at each timestep and *NPP* and *f N_up_* are averaged over a time period *τ_N_*, in a similar manner to environmental drivers. The first term in each equation represents the uptake capacity for either C or N given current environmental conditions, while the second term represents the amount already available to the plant. In addition to these terms, the N-limited growth *N_growth_*, includes an additional flux, the nutrients reabsorbed before leaf shedding. All variables are averaged over a period of *τ_N_*, as above.

Given the availability of C and N, the direction variable *D_N_*is then calculated as:

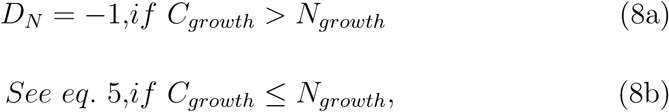

so that the leaf N content decreases if there is not sufficient N available for growth and defaults to the optimal C export variant if there is.

### 2.2 Optimality approach

For the purpose of the optimal C export and optimal growth model variants, we apply an optimality criteria which states that plants aim to maximise the canopy carbon export and growth respectively given moving averages over a period of *τ_N_* (30 days) of all environmental drivers as well as relevant model state variables, such as NPP and N uptake (Eq. 6 and 7). The optimality criteria is applied at every timestep, resulting in a continuously changing values of the direction variable *D_N_* and therefore a continuously changing leaf N value. The *τ_N_* parameter is meant to buffer sub-daily and abrupt changes in driving variables. The short timestep ensures that, although the approach only calculates the direction of change, the change in leaf N is smooth.

We use a numerical approach to solve the optimality problem for two reasons. The first is that, while using a TBM with an explicitly layered canopy and complex representation of photosynthesis produces more realistic predictions, it also means that the problem is non-linear and has no analytic solution. The second reason is that one of the central concepts of our approach is that we are not solving for the leaf N values that gives the actual maximum C export or growth at any point in time but rather assume that plants tend towards equilibrium, given physiological and biochemical constraints to their rate of change.

### 2.3 Site description

We test all model variants at two Free Air CO_2_ Enrichment (FACE) sites, the Duke Forest (hereafter Duke) and Oak Ridge National Laboratory (hereafter ORNL) experimental sites as well as at the Harvard Forest N addition experiment.

The Duke FACE experiment (McCarthy *et al*., 2007) was carried out in a loblolly pine (*Pinus taeda*) plantation (35.9*^◦^*N, 79.08*^◦^*W). The experiment began in August 1996 and consisted of three plots with ambient and three plots with elevated (+200 ppm) CO_2_ concentrations, paired according to soil N availability. The Oak Ridge FACE experiment (Norby *et al*., 2002) was set up in a sweetgum (*Liquidambar styraciflua L.*) plantation (35.9*^◦^*N, 84.33*^◦^*W) and begun in April 1998. It included five experimental plots, of which three were at ambient and two at elevated CO_2_ (average of 547 ppm) concentrations. For both sites we use annual measurements of leaf N content, leaf biomass and net primary productivity for model evaluation (Finzi *et al*., 2007; Walker *et al*., 2014; Zaehle *et al*., 2014). At both sites, the NPP is derived from direct and indirect measurements of leaf, fine root and wood growth.

The chronic nitrogen addition experiment (Magill *et al*., 2004) located at Harvard Forest (42.5*^◦^*N, 72.16*^◦^*W) is part of the Harvard Forest Longterm Ecological Research (LTER) site. The experiment consists of four plots each in a red pine (*(Pinus resinosa*) and a hardwood stand (dominated by black and red oak, *Quercus velutina Lam.; Q. rubra L.*)). Of the four plots in each stand, we used the control (no N addition), as well as the low N addition (5 gN m*^−^*^2^ yr*^−^*^1^), and high N addition (15 gN m*^−^*^2^ yr*^−^*^1^) treatments, excluding the nitrogen and sulphur treatment as this was shown to not be significantly different from the low N treatment and is also outside the scope of this paper. Nutrient additions started in spring of 1988 and were made in equal doses at 4-week intervals throughout the growing season, with the last application taking place in 2004. For model evaluation, we use woody biomass increment from Magill *et al*. (2004) and annual leaf N content from the Harvard Forest Data Archive (Frey S, 2018). For the detailed analysis we show only results from the hardwood stand, as observations for the pine stand shows a decrease in biomass with N addition, as this is a result of processes not relevant to the current analysis.

In addition to this detailed analysis, we also run the model at 16 additional N fertilisation sites located in Northern Europe and North America, including evergreen needleleaf and broadleaf deciduous forests. A number of these sites include plots with different levels of N addition, resulting in 23 separate experimental plots. The rate of N addition varies widely (1.7 - 15 gN m*^−^*^2^year*^−^*^1^) as does the duration of the experiment (1 - 40 years). We extract all biomass and leaf N response data from the respective experiment papers where available. See Table S2 for details for each site. For the purpose of this analysis, we exclude sites with a pronounced decrease in biomass under fertilisation, namely Asa, Norrliden N3 and Harvard pine.

### 2.4 Model protocol and input data

The QUINCY model requires as input half-hourly meteorological drivers (short-and longwave radiation, air temperature, precipitation (as rain and snowfall), air pressure, humidity and wind velocity), as well as atmospheric CO_2_ concentration and nitrogen and phosphorus deposition rates. Additional model inputs are vegetation type, and soil physical and chemical parameters (texture, bulk density, rooting and soil depth, as well as inorganic soil P content). Although the model has the option to include fully coupled N and P cycles, for the purpose of this paper we have used the CN-only model, where soil solute inorganic P availability is kept at a constant, non-limiting level.

The daily meteorological data for 1901 to 2015 was extracted from the CRUNCEP dataset, version 7, (Viovy, 2016), and disaggregated to the half-hourly model timestep using the statistical weather generator described in (Zaehle & Friend, 2010). The annually changing CO_2_ atmospheric concentration was taken from Le Quéré *et al*. (2018), and the time series of N deposition for each site from Lamarque *et al*. (2010) and Lamarque *et al*. (2011). In addition, for the two FACE sites, the local meteorological data as well as the CO_2_ concentrations for the duration of the experiments were used (Walker *et al*., 2014).

The soil and vegetation biogeochemical pools are brought to quasi-equilibrium through a 500 year model spinup, using forcing data from the period 1901-1930. We then run the model with transient climate and CO_2_ concentrations for the years 1901-2015 for all sites and treatments. Each site is harvested according to the harvest date available in the literature. After harvest, all biomass is retained in the system as litter, with the exception of the harvested fraction of the woody biomass (set to 80 %), which is removed. A side effect of harvesting is that as the four model variants have different growth rates, the sites can be at slightly different stages of succession at the start of each experiment.

For the N addition experiments, we introduce the additional N to the system to either or both the NH_4_ or NO_3_ soil inorganic pool depending on the chemical form of the fertiliser, in the quantity and with the timing described for each respective experiment.

As well as the above fully transient runs, we run the model to equilibrium for a range of soil N availabilities, to explore the behaviour of all model variants at equilibrium without the added behaviour of transient runs or experimental additions. For this purpose, we perform model runs using the meteorology and initial conditions of the control Harvard hardwood site. We run the model for 500 years with a fixed atmospheric CO_2_ concentration for a mature forest. We vary the available soil N level by varying the rate of biological N fixation (BNF). BNF in QUINCY is represented as an asymbiotic processes with a fixed, temperature-dependent rate given a soil mineral N threshold, so that effectively varying the rate of BNF results in varying levels of soil mineral N without affecting other model processes. We perform two sets of runs, with the CO_2_ concentrations set to 380 ppm and 550 ppm respectively.

### 2.5 Model parameterisation

We use the QUINCY model with its default parameterisation, without calibrating the parameters to any of the sites used in this study. Table 1 shows the parameters that differ between each of the four model variants and their values for the two PFTS used here, broadleaf deciduous forest and needleleaf evergreen forest. Table S1 lists other model parameters relevant to the four leaf N variants. For a full list of parameters see Thum *et al*. (2019).

Both the optimal C export and optimal growth introduce two new, PFT-independent parameters, in addition to those present in the standard QUINCY model, *δ_N_* and *τ_N_*. In comparison, the empirical model requires two PFT-specific parameters for leaf CN ratio bounds, two empirical parameters that drive the shape of the curve and the two parameters it has in common with the optimal variants (Table 1).

To test the model stability to variations in parameters, we perform a parameter sensitivity analysis, detailed in Section S1 of the Supplementary material 1.

## 3 Results

### 3.1 Model predictions at equilibrium

Figure 3 shows model NPP and canopy average foliar N content at equilibrium under a gradient of soil N availability for all four model variants, for ambient CO_2_ (380 ppm, (a) and (b)) and the relative response under elevated CO_2_ (550 ppm, (c) and (d)). This explores the theoretical response of the model at equilibrium, without transient climate or CO_2_ concentrations. This provides a prediction of a similar type to what previous optimal models have included, but derived from a fully prognostic carbon-nitrogen cycle model. It is also a clearer way of explaining the model runs at experimental sites below and testing the theoretical assumptions behind each model variant (Fig. 2).

**Figure 3:**
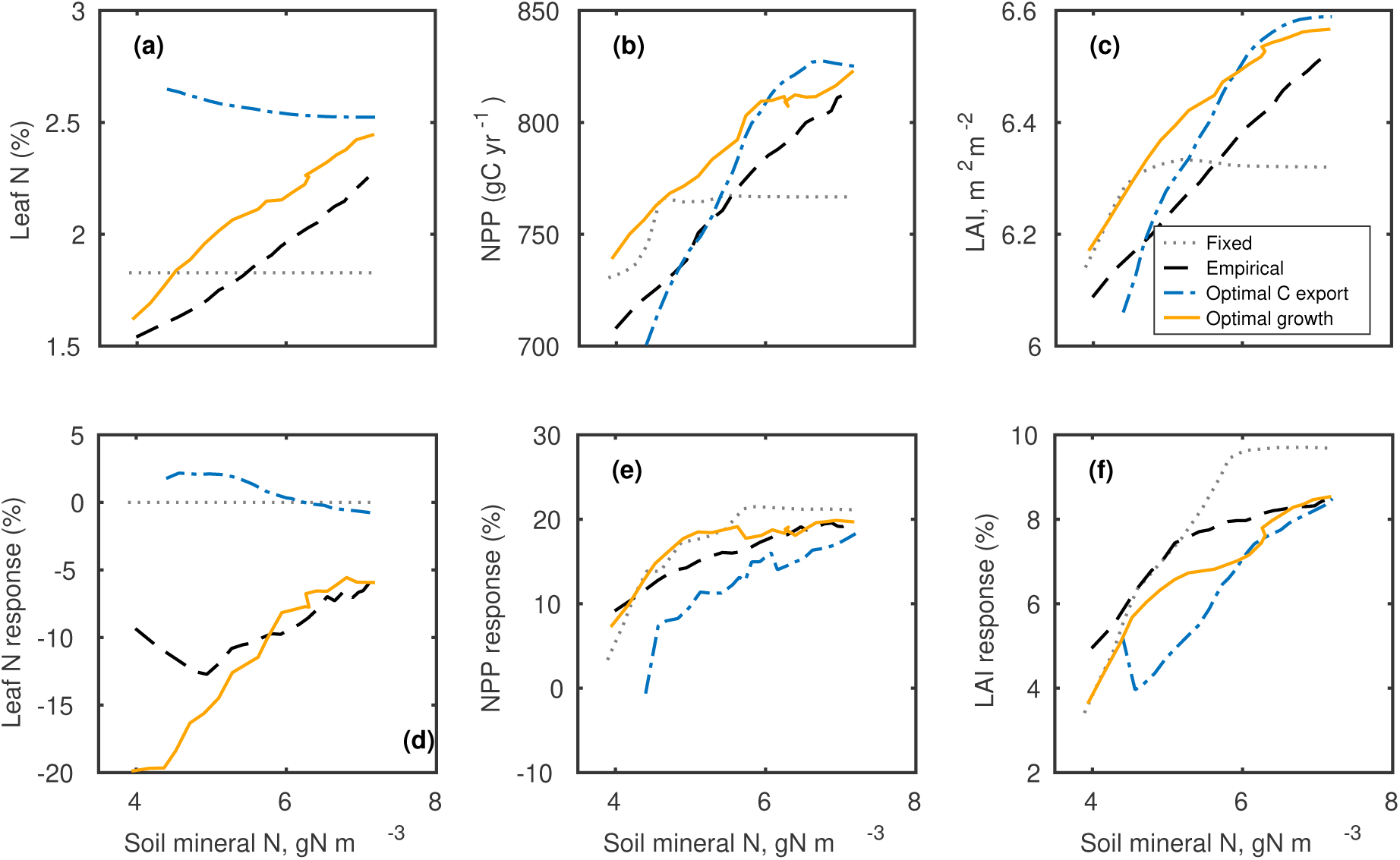
At-equilibrium whole-canopy leaf N content (a), NPP (b) and maximum annual LAI (c) over a range of available soil mineral N, as well as the response to elevated CO_2_ (550 ppm) relative to the predictions at 380 ppm ((d), (e) and (f)) for all model variants. Leaf N values are canopy averages. Leaf N values predicted by the optimal C export variant show a very strong positive response to elevated CO_2_ at very low availability and have been omitted for plot clarity. Soil N values refer to total mineral N to a depth of 1 m. All simulation are done using the meteorology and initial conditions of the control Harvard hardwood site, a temperate broadleaf deciduous forest.

Both the empirical and optimal growth model options predict an increase in leaf N content with increasing N availability, as expected, while the optimal C export model predicts a slight decrease, as it mainly responds to variations in LAI rather than plant N status. Generally, under lower N availability, the model predicts a lower LAI, meaning that there are no deeper, more shaded, canopy layers which would have higher respiration values compared to photosynthesis, thereby shifting the NPP response curve (Fig. S3 (a) and (b)) and increasing the optimal leaf N concentration. As there is a slight increase in LAI predicted by all model variants with an increase in soil N (Fig. 3(c)), there is a resulting slight increase in leaf N predicted by the optimal C export variant with increased N availability. This kind of response is also observed under increased CO_2_. At higher soil N values, when plants are not N-limited, the optimal C export and optimal growth versions produce similar predictions, as is expected from the assumptions of these two variants (Fig. 2, Eq. 8). All models with flexible leaf N show a higher NPP than the fixed variant at high soil N, however the empirical and optimal C export versions show lower NPP at low N availability. The optimal C export variant shows a leaf N response that does not match observations or our process understanding, namely the decrease in leaf N content with increased N availability (Fig. 3 (a)), and therefore a lower overall growth (Fig. 3(b)), demonstrating that a canopy C export only optimal approach does not produce physiologically realistic predictions. The optimal growth variant results in the highest NPP for most of the soil N range, as expected from the optimal criteria that maximises growth. At high soil N however, the optimal C export variant predicts a slightly higher NPP, as its higher N demand caused by the high leaf N content, can be met by the available soil N.

In terms of predictions under elevated CO_2_ (Fig. 3 (d) - (f)), both the empirical and optimal growth versions show a decrease in leaf N, strongest at low N availability, while the optimal C export shows only a very small change. The optimal C export model also shows an overall less pronounced response to elevated CO_2_ than the empirical and optimal growth versions.

### 3.2 Model response to elevated CO_2_

We show timeseries from fully transient model runs at the two FACE sites, including observed and predicted NPP and leaf N content (Fig. 4) as well as whole ecosystem responses as a mean of the first three years and the last three years of each experiment (Fig. 5). In terms of leaf N content, all model variants are able to predict the observed ambient values reasonably at both FACE sites, although the empirical option shows slightly higher values for Duke (Fig. 4 (a) and (b)). For the ORNL site, the optimal C export model overestimates observed values.

**Figure 4:**
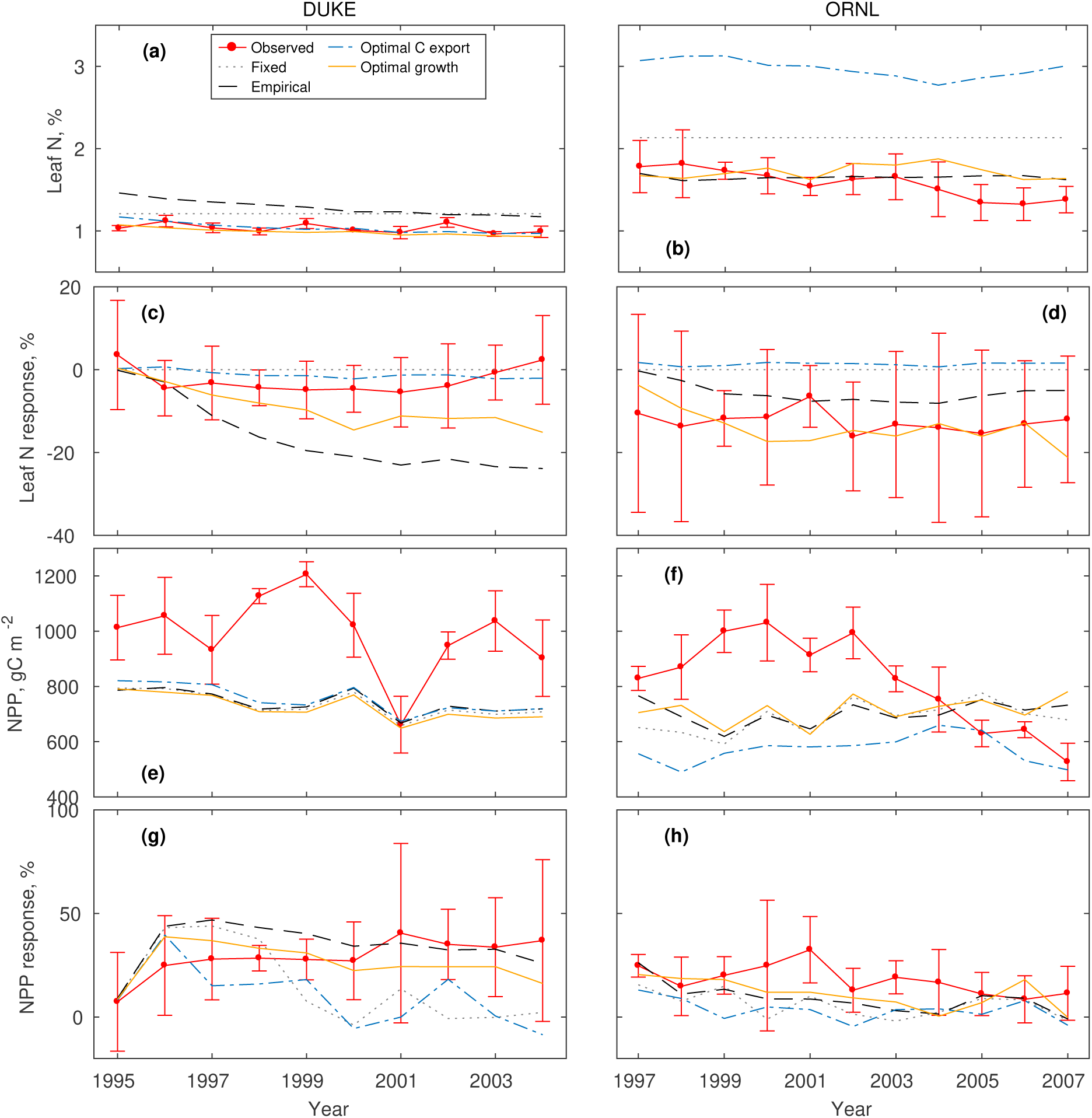
Model predictions at the Duke (left) and ORNL (right) FACE sites for absolute leaf N concentration ((a) and (b)), relative response of leaf N under elevated CO_2_ ((c) and (d)), absolute NPP values ((e) and (f)) and relative response of NPP ((g) and (h)).

**Figure 5:**
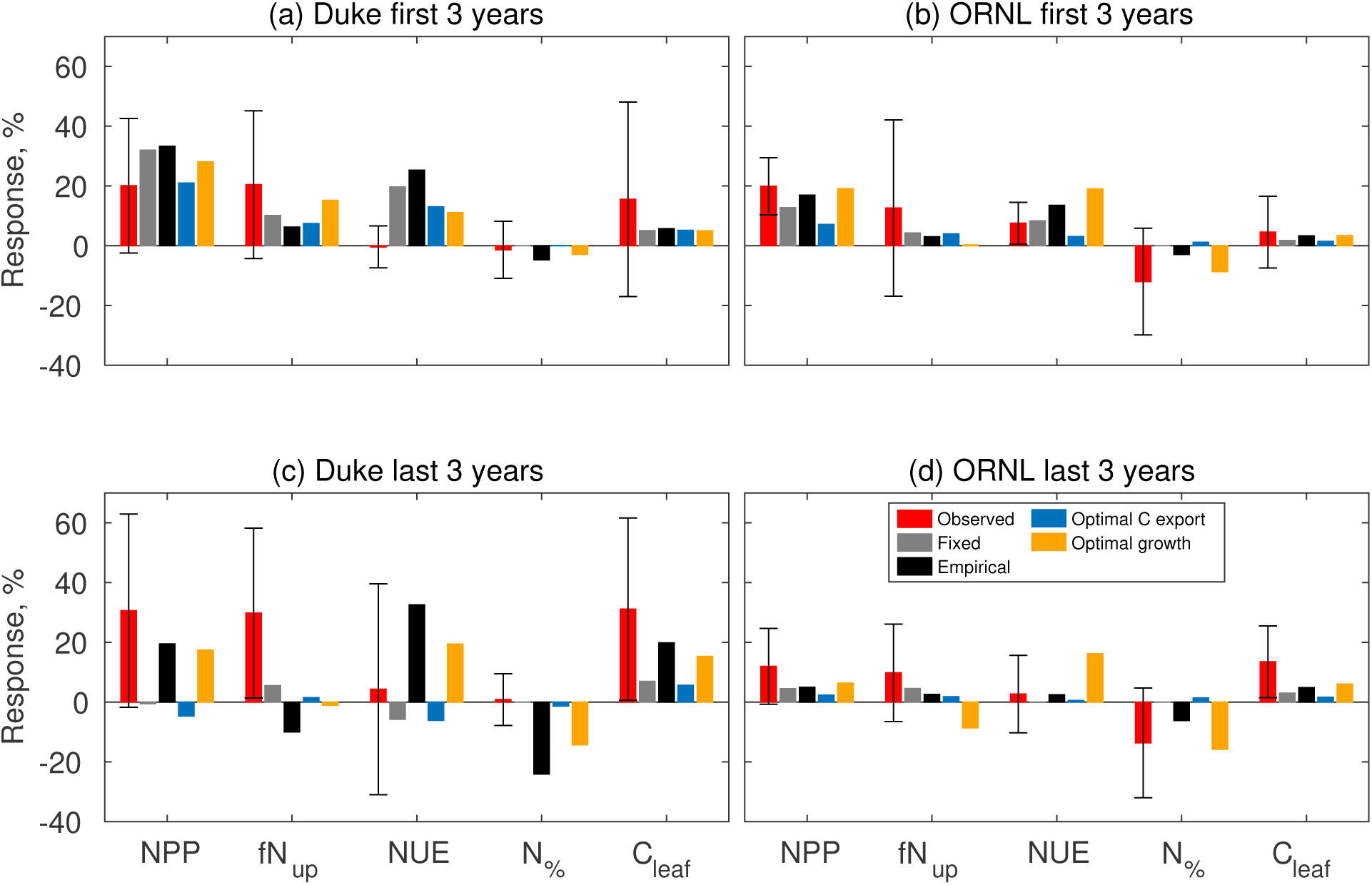
Ecosystem response to elevated CO_2_ at the Duke and ORNL FACE sites for the first 3 years ((a) and (b)) and last three years ((c) and (d)) for all model variants. Variables shown are net primary productivity (NPP), plant N uptake (*f N_up_*, nitrogen use efficiency (NUE), leaf N concentration (*N*_%_), maximum annual total canopy C (*C_leaf_*. All variables are shown as relative responses %.

Both the empirical and optimal growth model variants can correctly predict the direction of change in leaf N throughout the experiment under elevated CO_2_ for both sites (Fig. 4 (c) and (d)), while the optimal C export fails to show a significant change in leaf N concentrations under elevated CO_2_ (-1.3 % and 1.4 % for Duke and ORNL, respectively at the end of each experiment, Fig. 5(c) and (d)). For the Duke FACE both the empirical and optimal growth variants predict a decrease in leaf N; the observations also show a decrease (Fig. 4(c)), although not so pronounced and increasing slightly at the end of the experiment. While both model variants overestimate the magnitude of the change, the optimal growth does so to a lesser degree (-24.1% empirical and -14.3% optimal growth, Fig. 5(c)). In the case of the ORNL site, the optimal growth variant gives the prediction closest to observations (observed -13.6 %, empirical -6.1 %, optimal growth -15.8 %, Fig. 5(d)).

All model variants predict similar NPP at ambient CO_2_, generally underestimating observed values at both sites (Fig. 4(e) and (f)). The optimal C export variant predicts an even lower NPP at ORNL, caused by the predicted high leaf N value, which leads to a higher growth demand for N and therefore a lower resulting growth.

Observations at both the Duke and ORNL sites show a positive NPP response to elevated CO_2_ throughout the experiment (Fig. 4 (g) and (h)), although for ORNL this response decreases significantly towards the end of the experiment. The first notable feature is that all model variants are capable of predicting this positive response at the start of the experiment (Fig. 5(a) and (b)), although this is overestimated for all variants in the case of Duke. However, by the end of the experiment (Fig. 5 (c) and (d)), only the empirical and optimal growth are able to predict the sustained positive response for the Duke site (observed 30.6 %, empirical 19.5 %, optimal growth 17.4 %). All models predict a lower NPP response than expected for the entire duration of the ORNL experiment (observed 12.0 %, empirical 4.5 %, optimal growth 6.3 % at the end of the experiment, Fig. 5(d)).

All model variants predict a lower response in total canopy C at both sites (Fig. 5(c) and (d)), indicating a missing shift in biomass allocation. While there is an observed positive response in N uptake at both sites, all model variants predict a lower response, including a decrease in uptake in the case of the empirical variant at Duke and the optimal growth at ORNL at the end of the experiment (Fig. 5 (c) and (d)). Overall, we find that the optimal C export variant does not reproduce observed responses to elevated CO_2_ in either leaf N content or NPP, while the empirical and optimal growth variants both reproduce the direction of response correctly throughout both experiments, but the optimal growth captures better the magnitude of the observations.

### 3.3 Model response to N addition

In the case of the Harvard Forest N addition experiment (Fig. 7 and timeseries in Fig. 6), the observations indicate an increase in leaf N content with N addition (low N 3.3 %, high N 21.0 %), as do the empirical variant (low N 50.9 %, high N 64.0 %)) and optimal growth variant (low N 34.3 % and high N 45.4 %), even though they both overestimate this response (Fig. 7(c) and (d)). Notably, the optimal C export variant does not predict an increase in leaf N (low N -4.4 %, high N -4.3 % at the end of the experiment), as it responds to changes in LAI and not N availability, and the model predicts a low change in LAI and therefore a corresponding low response in leaf N (Fig. 7(c) and (d)). This is expected from the model assumptions (Fig. 2) and discussed for the at-equilibrium simulations. The peak in the leaf N response of the optimal growth variant (Fig. 6) is caused by a decrease in the control leaf N (but not the treatments), which coincides with the 2001-2002 drought; as there is a gap in the data in this period it is difficult to assess whether this reflects a real plant response.

**Figure 6:**
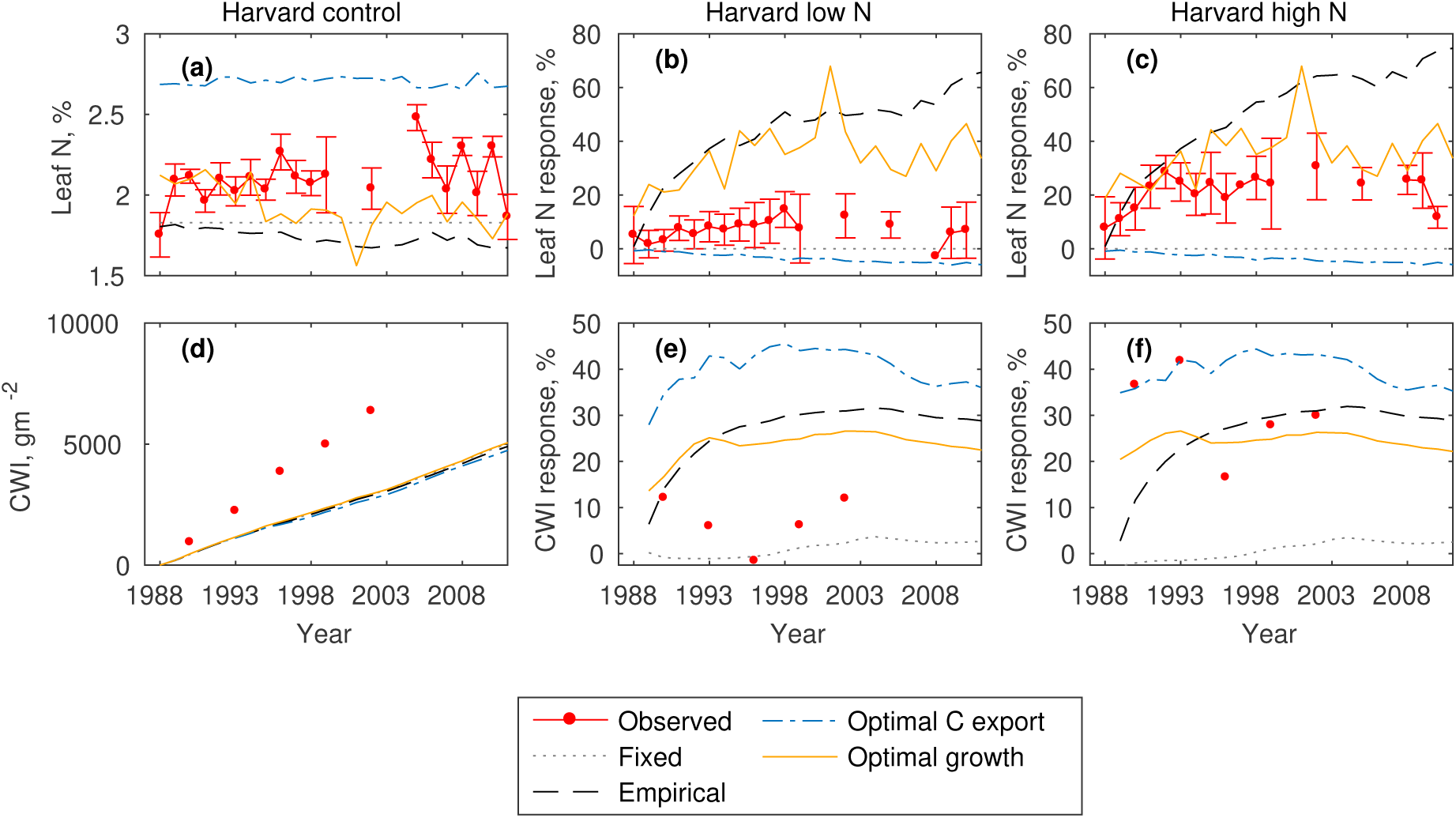
Model predictions at the Harvard long term N addition experimental site for absolute leaf N content values ((a)), relative response of leaf N under low ((b)) and high ((c)) N addition levels, as well as absolute current wood increment (CWI) ((d)) and relative response of CWI ((e) and (f)).

**Figure 7:**
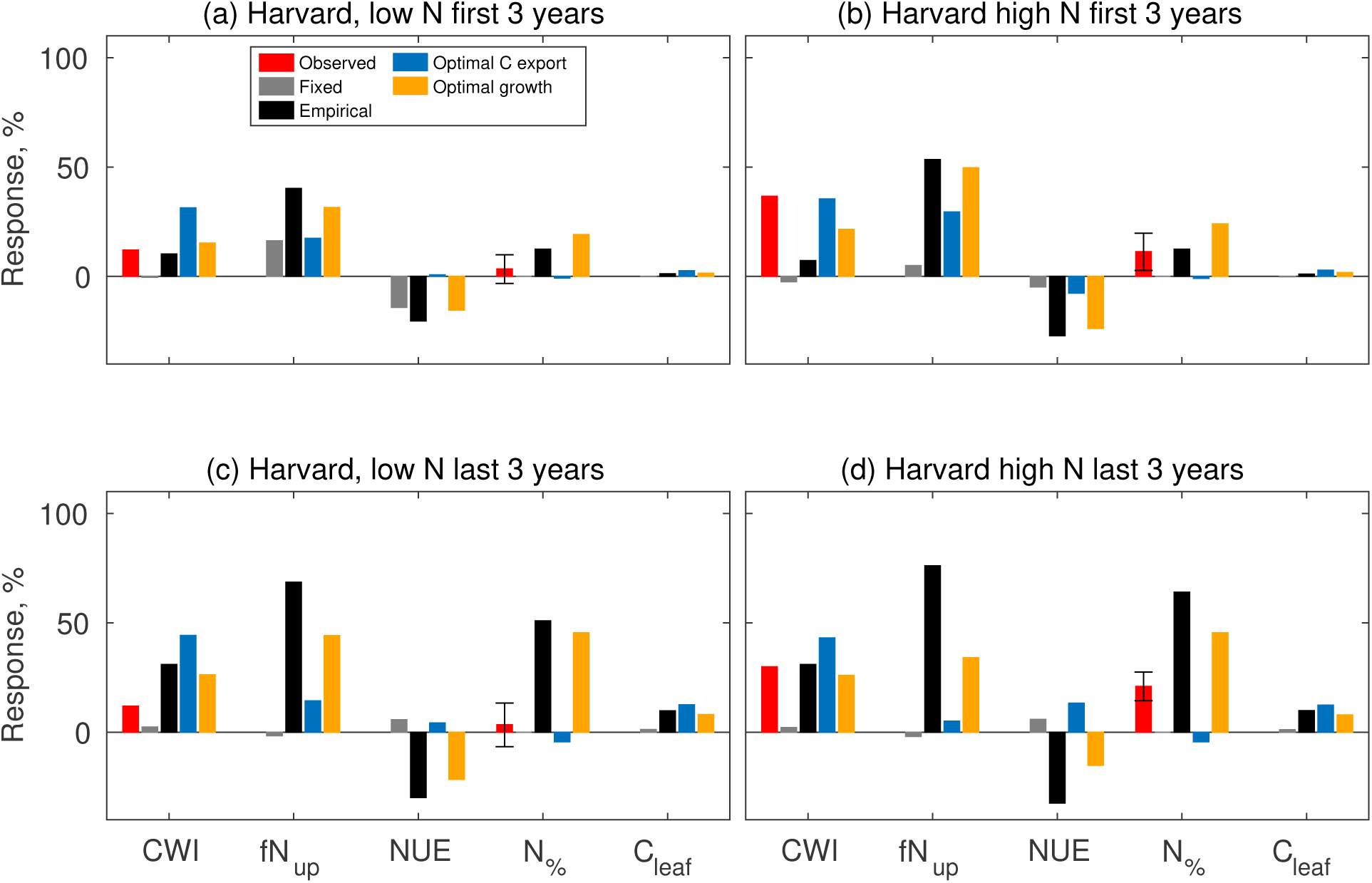
Ecosystem response to N addition at the Harvard hardwood site for the first 3 years ((a) and (b)) and last three years ((c) and (d)) for all model variants. Variables shown are current wood increment (CWI), plant N uptake (*f N_up_*, nitrogen use efficiency (NUE), leaf N concentration (*N*_%_), maximum annual total canopy C (*C_leaf_*. All variables are shown as relative responses %.

All model variants with flexible leaf N content predict a positive current wood increment (CWI) response both at the start and end of the experiment (Fig. 6(c) and (d)), although they all overestimate the response in the case of low N addition at the end of the experiment (observed 11.8 %, empirical 31.0 %, optimal growth 26.2 %, Fig. 7(c)). However, they show a better fit for the high N addition (observed 29.9 %, empirical 30.9 %, optimal growth 26.0 %, Fig. 7(d)). All variants underestimate the magnitude of observed CWI for the control plot (Fig. 6(d)).

To test the generality of the findings at the Harvard Forest site, we run our model for a selection of forest N fertilisation sites (Fig. 8). The magnitude of the predicted growth response for both the empirical and optimal growth model variants is linked to the average ambient temperature of the site, with a stronger response at colder sites (Fig. 8(a) and (b)). This is because the soil N availability, and therefore plant N limitation status is strongly dependent on temperature. The temperature dependency of plant response to N addition is present in reality, however this relation between observations and temperature is less evident than in the case of the model. This is caused by other confounding factors that drive local N availability in reality at each site, such as soil type and N deposition rates, factors which are not necessarily present in the model. The empirical variant largely reproduces observed biomass responses for warmer sites but over-estimates responses for the colder sites (Fig. 8(a)). On the other hand, the optimal growth variant has a tendency to underestimate the biomass response, especially for the higher observed responses (Fig. 8(a)). It is worth noting that the site with a very high observed biomass response (127 %) is one of the very northern sites, with very low annual average temperatures. Both variants show a better fit for the biomass response for broadleaved deciduous forests (normalised root mean squared error, NRMSE = 0.49 empirical and 0.55 optimal growth) than for the needleleaved evergreen (NRMSE = 1.88 empirical and 1.82 optimal growth). For the leaf N content observations (Fig. 8(c) and (d)), both model variants perform similarly, with the empirical one tending to predict a higher leaf N response. Unlike the biomass response there is no relation between ambient temperature and leaf N in either the model or the data. The optimal growth variant shows a better fit for the broadleaved site leaf N response than the empirical, although it is worth noting that the only deciduous site with leaf N measurements is Harvard Forest (discussed in detail above) so it is difficult to generalise this finding. For the evergreen forest sites, both the empirical and optimal growth variants perform similarly (NRMSE of 0.95 and 1.10 respectively). The two data points with very high leaf N response, which cannot be reproduced by either variant, are both the same site with different N addition levels and as McNulty *et al*. (2005) note, the observed values are unusually high.

**Figure 8:**
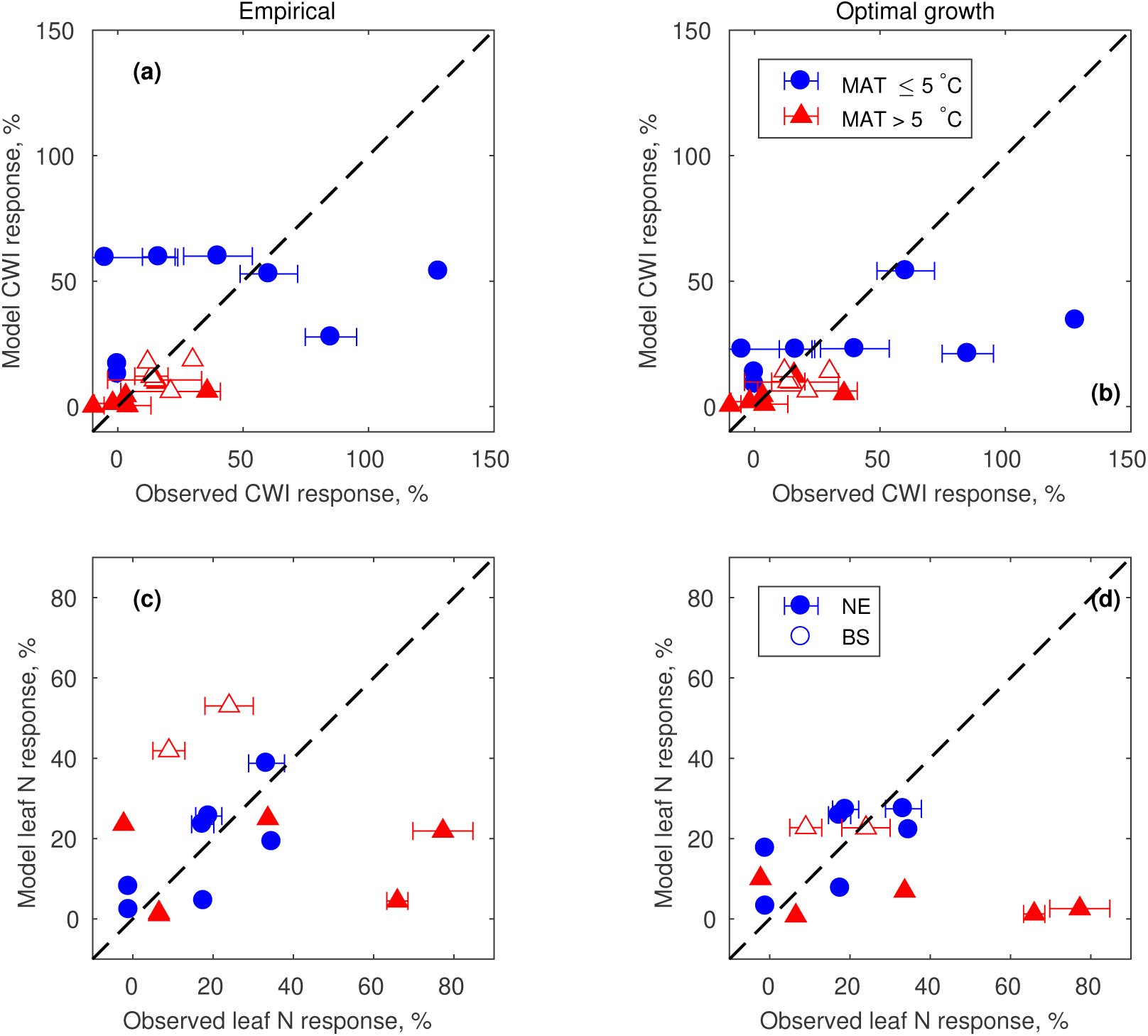
Biomass ((a) and (b)) and leaf N ((c) and (d)) response for a set of forest N addition sites as predicted by the empirical (left) and optimal growth (right) model variants, grouped by PFT (BS - broadleaved seasonal, open symbols, NE - needleelaved evergreen, closed symbols) and average annual temperature (mean annual temperature ¡ 5*^◦^*, blue circles and mean annual temperature ¿ 5*^◦^*, red triangles.

A complete uncertainty analysis is beyond the scope of the paper, however, we have included a parameter sensitivity study for two of the sites used in this study, Duke FACE and Harvard hardwood high N, for the empirical and optimal growth variants (Fig. S1 and S2), which shows that model predictions are stable and robust. Model uncertainty in absolute values of leaf N content, NPP and biomass production is similar for both variants. The uncertainty in the response under elevated CO_2_ is largely reduced for the optimal growth variant, although for the N addition site both variants show a high uncertainty.

## 4 Discussion

### 4.1 Model performance

In the current paper, we test the hypothesis that changes in leaf nitrogen content can be represented in a terrestrial biosphere model using the optimality principle, by assuming that leaf N changes dynamically in order to maximise plant growth, from the perspective of both net canopy C export and whole plant nutrient status. We show that an optimal scheme which focuses solely on the plant C balance can not represent plant responses at the two FACE and one N fertilisation site we used. Additionally, the empirical formulation, while performing better than the optimal C export variant under both elevated CO_2_ and N fertilisation, overestimates the magnitude of changes in leaf N (Fig. 4 and 6).

All four model variants underestimate total biomass growth, as either NPP for the two FACE sites (Fig. 4) or current wood increment for the Harvard forest site (Fig. 6). The predicted absolute biomass values are strongly dependent on initial conditions and model spinup, specifically on the soil nutrient status. In the current paper we use the default QUINCY soil model and we have not attempted to improve its representation, therefore the present model biases are intrinsic to its structure, but common to all leaf N model variants. The soil initial condition and process representation is also why the model, while representing the direction of change in leaf N under N addition, largely fails to represent the temporal trend (Fig. 6(b) and (c)).

It is important to note that the model has not been calibrated to any of the sites used in this study. In fact, one of the advantages of the optimality approach is that the property considered optimal, in our case leaf N content, becomes an emergent property of the model. This means that optimal models are more general and portable across sites and ecosystems.

### 4.2 Implications of the optimality approach

The majority of previous studies implement optimality as an at-equilibrium process, meaning that plants have the capacity to reach the optimum at any given time. This is done either in a model that only predicts steady-state vegetation processes or, when incorporated into a TBM (e.g. Ali *et al*., 2015), the plant property being optimised is set to its optimum value at each timestep. In the current study, we choose a ‘towards equilibrium’ approach, in that the optimality criterion only gives the direction of change and the magnitude of the actual change is limited to a given amount, meant to mirror physiological and ecological limitations to plant plasticity (Valladares *et al*., 2007). Therefore, this optimality approach can be used to simulate transient plant acclimation under variable environmental conditions at the timescale that a TBM is usually run at, whilst at the same time maintaining biologically realistic change rates of leaf N concentrations.

Another novel aspect of the optimality scheme we present here is the inclusion of a whole-plant optimality criterion, in contrast with existing studies which have focused on canopy C assimilation or export (e.g. Dewar, 1996; Ali *et al*., 2015; Smith *et al*., 2019). We show that a carbon only optimality criterion can not represent observed plant responses at the three main experimental sites we used. As shown in Fig. 3, the optimal C export variant varies the leaf N content in response to changes in LAI and environmental conditions that drive variation in photosynthesis, rather than directly changes in N availability. In particular, it predicts an increase or no change in leaf N under elevated CO_2_, as the photosynthesis rate per unit N increases, but respiration remains constant, a behaviour we see both in theory (Fig. 2) and in at-equilibrium and transient simulations. This has implications for productivity estimates and therefore for predictions under future conditions, as seen at both FACE sites (Fig. 4) in this study, where the optimal C export variant cannot predict the positive response of NPP over the duration of the experiment as well as at the N addition site (Fig. 6), where it predicts a too high growth response. Our results highlight the importance of including whole-plant responses in order to correctly represent growth, especially when taking into account nutrient limitation.

A previous model inter-comparison at FACE sites (Zaehle *et al*., 2014) has shown that models that include an empirical variation in leaf N content tend to overestimate the decrease in leaf N, something which this study also shows, specifically at the Duke site (Fig. 4(c)). The optimal growth model also predicts a too strong response in leaf N, both at the Duke FACE and the Harvard N addition sites but to a lesser extent. From the theoretical, at-equilibrium results shown in Fig. 3 we can see that the differences in response to elevated CO_2_ between the empirical and the optimal growth vary with soil N availability so that it is possible that the mismatch between the observations and the optimal growth scheme is caused by the wrong initial soil conditions, as is also indicated by the too low predicted NPP and wood growth at both the FACE and the Harvard sites. It is important to note that the empirical functional form used here includes upper and lower bounds to leaf N variation derived from observations, while the optimal formulation emerges from the processes and interactions already included in the model. This means that the optimal approach has less empirical, PFT-specific parameters and relies only on our understanding of plant physiological processes. The optimal growth variant allows for plastic short-term response to environmental conditions, without the need for a change in PFT and is therefore a method more generally applicable in space and time.

Beyond the choice of optimal model there is of course also the choice of empirical formulation. Here, we use the formulation of (Zaehle & Friend, 2010), as implemented in QUINCY (Thum *et al*., 2019). It aims to constrain leaf N values within observed ranges and responds to whole-plant nutrient limitation. Other TBMs use similar response functions, making it a good benchmark for existing representations of leaf N versus optimal representations.

### 4.3 Extending the optimality approach to other model processes

The optimality approach we present here focuses solely on the representation of dynamic leaf N content, while keeping the baseline model structure of QUINCY as described in Thum *et al*. (2019). Dynamic variation in leaf N content clearly plays a key role in model predictions of ecosystem productivity, as shown here, a number of other key canopy processes do so also, as discussed below. Optimality concepts have been proposed as a way to represent many of these processes and future development of our model can incorporate such representations.

Changes in atmospheric CO_2_ have been shown to lead to changes in the photosynthetic apparatus, a process which is often represented mathematically as a shift in the allocation of photosynthetic N between the different components to maximise C assimilation at the leaf level, a representation sometimes referred to as the coordination hypothesis (Medlyn, 1996; Ali *et al*., 2015; Smith *et al*., 2019). The inclusion of this hypothesis within our optimal leaf N model would modify the shape of the NPP response to leaf N content (Fig. 2) thereby changing the optimal leaf N under given conditions, especially for the optimal C export model variant potentially leading to a more pronounced response under elevated CO_2_.

Here, we assume that carbon export and plant nutrient status are the only drivers of changes in leaf N. While we can consider this assumption to hold at longer than annual timescales, at seasonal scales two more processes come into play: leaf ageing and N distribution with depth in the canopy. The QUINCY model assumes a constant C:N ratio throughout a leaf’s lifetime, but in reality leaf N decreases with leaf age (Reich *et al*., 1991; Kitajima *et al*., 1997). This process is particularly important in evergreen species, and has been shown to lead to changes in ecosystem productivity in tropical forests (Caldararu *et al*., 2012; Wu *et al*., 2016). Combining an age-based decline in leaf N with our optimality model would provide further constraints on the flexibility of leaf N values, potentially reducing the too-large response currently predicted.

We represent the distribution of N in the canopy as an exponentially decreasing function with depth (and therefore light level), with parameters based on observations (Zaehle & Friend, 2010), which is consistent with numerous observational studies (e.g. Meir *et al*., 2002; Kull & Niinemets, 1998). However, this distribution is time invariant, although we would expect it to vary with changes in the light environment in time and space (Meir *et al*., 2002). Previous studies (e.g. Dewar *et al*., 2012) have used the optimality approach to also predict the within-canopy N distribution albeit at steady state and it should therefore be possible in future versions of our model to include this further flexible process.

Leaf N content is known to co-vary with specific leaf area (SLA) (e.g. Wright *et al*., 2004), a process which we have not included in the current version of QUINCY. While the co-variation is an established fact, the actual causes of variation in SLA are complex, including structural and biochemical drivers (Poorter *et al*., 2009). Evans & Poorter (2001) present an optimality-based model which explores the trade-offs between N allocation and changes in leaf structure and SLA, which can be a way forward for a flexible and realistic way of representing variations in SLA. A flexible SLA value in QUINCY would affect the response of the C export variant as this depends strongly on LAI (Fig. 3), which will vary strongly with changes in SLA.

Our results highlight a number of key discrepancies in model predictions. All model variants fail to represent the observed increase in both nitrogen uptake and leaf biomass at the two FACE sites (Fig. 5), pointing to a need for dynamic biomass allocation. Plant flexible biomass allocation in response to nutrient and water availability is a well documented process (Poorter & Nagel, 2000; Hermans *et al*., 2006) and one that has previously been shown to impact models’ predictive capability under elevated CO_2_ (De Kauwe *et al*., 2014). In the current study, for clarity and to avoid complex model feedbacks, we have chosen to keep biomass allocation as a purely allometric process, independent of plant nutrient demand. The current model version has the option to include an empirical dynamic biomass allocation, following Zaehle & Friend (2010), but more process-based ways of representing this are also available including a scheme for optimal allocation (Mäkelä *et al*., 2008; Franklin, 2007). Such approaches would represent an increase in belowground allocation in response to nutrient and water limitation. A representation of dynamic allocation would lead to more N being available to plants, through an increase in overall N uptake, which would lead to a less pronounced decrease in leaf N as predicted by both the empirical and optimal growth variants, specifically at the Duke FACE site, and therefore a more accurate prediction. Similarly, under N addition, a flexible allocation scheme would reduce the fraction of biomass allocated to roots and therefore reduce N uptake, leading to a less pronounced increase in leaf N content, which is too high for both the empirical and optimal growth variants. This in turn would lead to lower C assimilation rates and therefore a reduction in the growth response, which is currently too high for all model variants.

### 4.4 Concluding remarks

In conclusion, we show that a whole plant optimality approach incorporated into a TBM can reproduce observed NPP and leaf N content responses for both elevated CO_2_ and N addition experiments and that an optimal approach which considers carbon export only cannot reproduce these responses. While both an empirical and a whole-plant optimality approach capture observations, with their own particular biases, we argue for the use of the optimality approach as being more rooted in physiological and ecological concepts. Our study shows how optimality concepts can be implemented in terrestrial biosphere models, making use of existing variables and parameters, to reproduce plant plastic responses in a biologically realistic manner.

## Acknowledgements

This work was supported by the European Research Council (ERC) under the European Union’s Horizon 2020 research and innovation programme (QUINCY; grant no. 647204). LY was supported by the framework of Priority Program SPP 1685 “Ecosystem Nutrition: Forest Strategies for Limited Phosphorus Resources” of the German Research Foundation (DFG), grant No.ZA 763/2-1. We are grateful to our scientific programmer, Dr. Jan Engel, for technical assistance in developing the code.

## Author contributions

SC and SZ designed the study and performed the analyses. SC, TT, LY and SZ developed the model. All authors contributed to writing the manuscript.

## Data availability

The FACE experiment data is freely available through the FACE Data Management System https://facedata.ornl.gov/. The Harvard Forest chronic nitrogen addition experiment foliar N data is freely available from the Harvard Forest Data Archive http://harvardforest.fas.harvard.edu:8080/exist/apps/datasets/showData.html?id=hf008. Compiled literature data on biomass and leaf N responses at N fertilisation experimental sites can be found as Supplementary material 2.

## Code availability

The scientific part of the code is available under a GPL v3 licence. The scientific code of QUINCY relies on software infrastructure from the MPI-ESM environment, which is subject to the MPI-M-Software-License-Agreement in its most recent form (http://www.mpimet.mpg.de/en/science/models/license). The source code is available online (https://git.bgc-jena.mpg.de/quincy/quincy-model-releases), but its access is restricted to registered users. Readers interested in running the model should request a username and password from the corresponding authors or via the git-repository. Model users are strongly encouraged to follow the fair-use policy stated on https://www.bgc-jena.mpg.de/bgi/index.php/Projects/QUINCYModel.

