## Supplementary figures and tables for "Whole-plant optimality predicts changes in leaf nitrogen under variable CO_2_ and nutrient availability"

### S1 Model sensitivity analysis

We use latin hypercube sampling (LHS) to test the model's sensitivity to its parameterisation and the robustness of the model structure (Saltelli *et al.*, 2000; Zaehle *et al.*, 2005). As the QUINCY model has a large number of parameters, we select a subset directly relevant to changes in leaf N for a total of 23 parameters (see Table 1 and S1). We use the LHS to sample 500 sets of uncorrelated parameters, each taken from a uniform distribution around 90% to 110% of its default value. We then run the model for the empirical and optimal growth variants with these parameter sets and use the resulting distribution of model output to calculate the mean and 95th percentile confidence intervals for variables of interest.

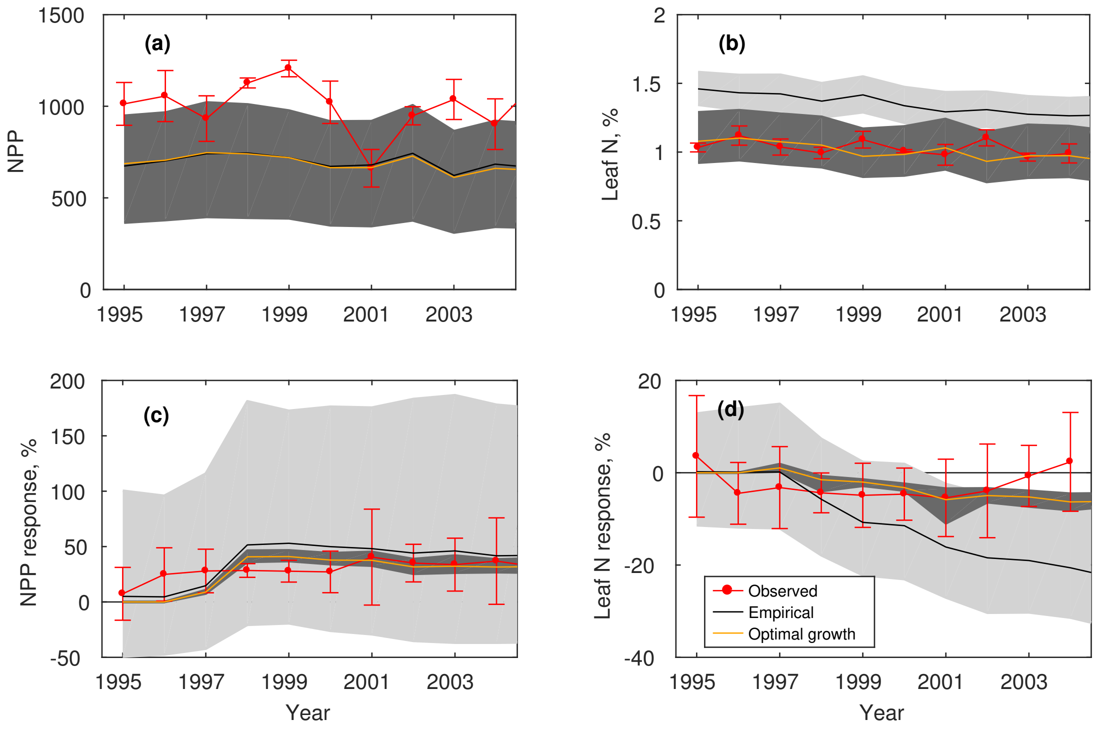


Figure S1: Timeseries of net primary production, NPP and leaf N content, with model uncertainty bounds for the empirical (black, light gray shading) and optimal growth (orange, dark grey shading) variants at the Duke FACE site.

To test the stability of the model given the baseline parameterisation, we show model uncertainty for absolute and relative NPP as well as leaf N content for the empirical and optimal growth variants (Fig. S2 and S1). Both variants result in realistic values and stable model output for both NPP and leaf N absolute values and relative responses for the parameter range tested.

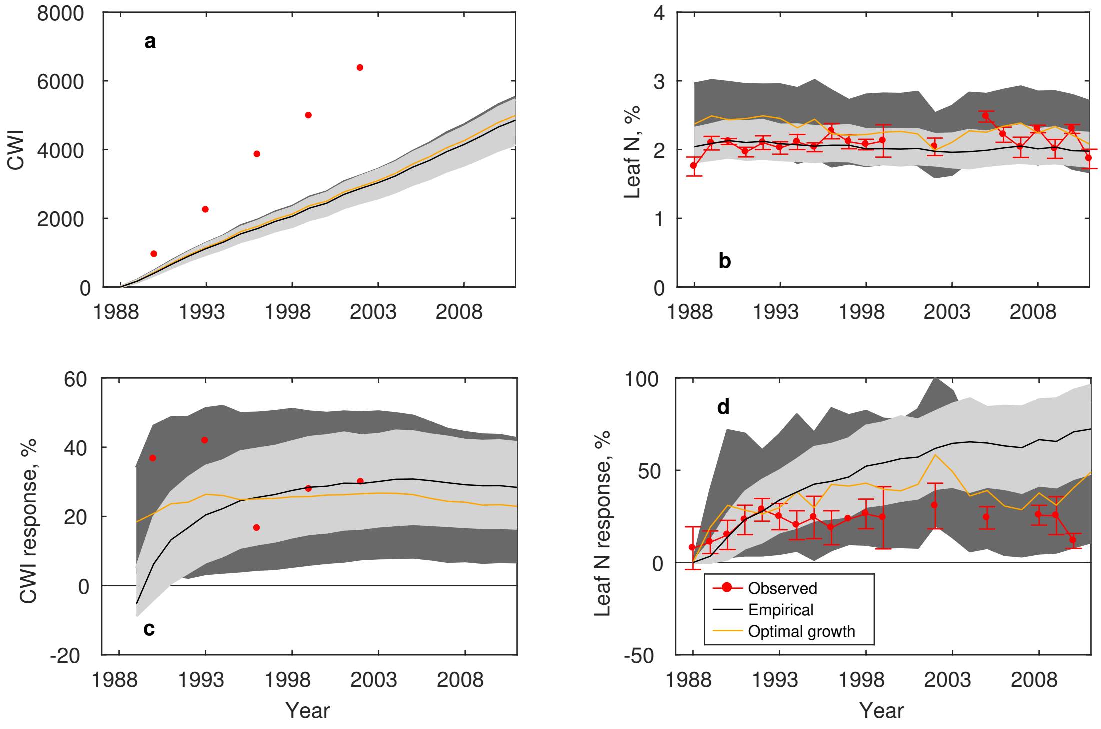


Figure S2: Timeseries of current woody increment, CWI and leaf N content, with model uncertainty bounds for the empirical (black, light gray shading) and optimal growth (orange, dark grey shading) variants at the Harvard Forest N addition site for the high N addition plots.

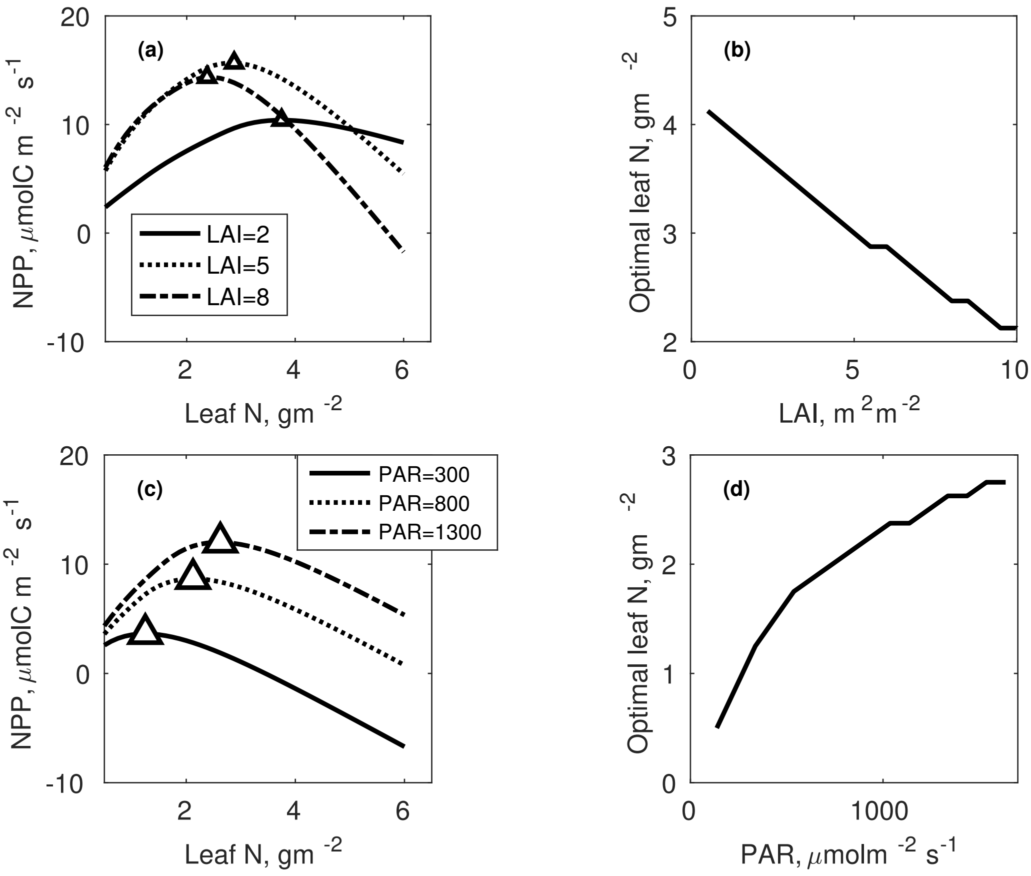


Figure S3: Theoretical representation of the response of optimal leaf N as predicted by the optimal C export variant in response to variation in LAI ((a) and (b)) and incident photosynthetically active radiation, PAR ((c) and (d)). Left-side panels show the dependency of net C assimilation and Leaf N for three different LAI values ((a)) and three different PAR values ((c)). Triangles show the leaf N for which maximum NPP occurs, equivalent to the optimal C export variant. Right-side panels show the resulting optimal leaf N variation with LAI and PAR, respectively.

Table S1: QUINCY model parameters relevant to the four formulations of dynamic leaf N. Note that this is not an exhaustive list of the model parameters and the full list can be found in (Thum *et al.*, 2019). BS = broadleaf deciduous, NE = needleleaf evergreen

| Symbol | Description | Value BS | Value NE | Unit | Citation |
| --- | --- | --- | --- | --- | --- |
| $f_{resp,growth}$ | Growth respiration fraction per unit new biomass | 0.25 | 0.25 | $\frac{molC}{molC}$ | Sprugel *et al.* (1995) |
| $f_{resp,maint}$ | Maintenance respiration rate for fine roots and leaves | 1.0 | 1.0 | $\frac{\mu molCO_2}{mmolN\ s}$ | Sprugel *et al.* (1995) |
| $r_{J2V}$ | Ratio of Jmax25/Vcmax25 | 1.97 | 1.97 | - | Wullschleger (1993) |
| $k_0^{chl}$ | Chlorophyll distribution with canopy depth | 6.0 | 6.0 | - | Zaehle & Friend (2010) |
| $k_1^{chl}$ | Chlorophyll distribution with canopy depth | 3.6 | 3.6 | - | Zaehle & Friend (2010) |
| $k_{fn}^{chl}$ | Chlorophyll distribution with canopy depth | 0.7 | 0.7 | - | Friend (2001) |
| $k_1^{struc}$ | Slope of structural leaf N with total N | $7.14 \times 10^3$ | $7.14 \times 10^3$ | $g^{-1}N$ | Evans (1989) |
| $k_{resorb}^{leaf}$ | Fraction of nutrient resorption before leaf shedding | 0.5 | 0.5 | - | Thum *et al.* (2019) |
| $k_{resorb}^{wood}$ | Fraction of nutrient resorption before wood death | 0.2 | 0.2 | - | Thum *et al.* (2019) |

Table S1 ctnd.: QUINCY model parameters relevant to the four formulations of dynamic leaf N. Note that this is not an exhaustive list of the model parameters and the full list can be found in (Thum *et al.*, 2019). BS = broadleaf deciduous, NE = needleleaf evergreen

| Symbol | Description | Value BS | Value NE | Unit | Citation |
|---|---|---|---|---|---|
| $\chi_{root}^{C:N}$ | Relative C:N of fine roots compared to leaves | 0.85 | 0.85 | - | Zaehle & Friend (2010) |
| $\chi_{wood}^{C:N}$ | Relative C:N of woody biomass compared to leaves | 0.145 | 0.145 | - | Zaehle & Friend (2010) |
| $K_{demand}^{half,N}$ | Fraction of target labile N at which uptake is reduced to 50% | 0.75 | 0.75 | - | Thum *et al.* (2019) |
| $k_{demand}$ | Nutrient uptake response function to labile nutrient concentration | 2.0 | 2.0 | - | Thum *et al.* (2019) |
| $k_0^{struc}$ | Maximum fraction of structural foliar N | 0.63 | 0.83 | - | Friend *et al.* (1997); Kattge *et al.* (2011) |
| $fN_{struc,cl}^{min}$ | Minimum fraction of structural foliar N | 0.45 | 0.65 | - | Thum *et al.* (2019) |
| $\tau_{leaf}$ | Turnover time of leaves | 0.48 | 3.31 | years | Kattge *et al.* (2011) |
| $\tau_{fine\_root}$ | Turnover time of fine roots | 0.7 | 0.7 | years | Ahrens *et al.* (2014) |
| $\tau_{growth}$ | Response timescale for processes related to plant growth | 1 | 1 | years | Thum *et al.* (2019) |

| Site | Coordinates | PFT | Duration | Annual N | Total N | Reference |
|---|---|---|---|---|---|---|
| Harvard hardwood low N | 42°30′N/72°10′W | BS | 16 | 5.0 | 83.8 | Magill *et al.* (2004) |
| Harvard hardwood high N | 42°30′N/72°10′W | BS | 16 | 15.0 | 251.3 | Magill *et al.* (2004) |
| Harvard pine low N | 42°30′N/72°10′W | NE | 16 | 5.0 | 83.8 | Magill *et al.* (2004) |
| Harvard pine high N | 42°30′N/72°10′W | NE | 16 | 15.0 | 251.3 | Magill *et al.* (2004) |
| Lake Laflamme low N | 47°17′N/71°14′W | NE | 3 | 1.7 | 5.1 | Houle & Moore (2008) |
| Lake Laflamme high N | 47°17′N/71°14′W | NE | 3 | 3.0 | 9.0 | Houle & Moore (2008) |
| Mt. Ascutney low N | 43°26′N/72°27′W | NE | 14 | 1.6 | 22.0 | McNulty *et al.* (2005) |
| Mt. Ascutney high N | 43°26′N/72°27′W | NE | 14 | 3.1 | 44.0 | McNulty *et al.* (2005) |
| Michigan B | 45°33′N/84°51′W | BS | 8 | 3.0 | 24.0 | Pregitzer *et al.* (2004); Pregitzer & Burton (2008) |
| Michigan C | 44°23′N/85°50′W | BS | 8 | 3.0 | 24.0 | Pregitzer *et al.* (2004); Pregitzer & Burton (2008) |
| Michigan D | 43°40′N/86°09′W | BS | 8 | 3.0 | 24.0 | Pregitzer *et al.* (2004); Pregitzer & Burton (2008) |
| Sodankylä | 67°42′N/26°13′E | NE | 27 | 2.8 | 75.0 | Andersson *et al.* (1998); Hyvönen *et al.* (2008) |
| Norråker | 64°27′N/15°34′E | NE | 24 | 4.0 | 96.3 | Andersson *et al.* (1998); Hyvönen *et al.* (2008) |
| Åseda | 57°06′N/15°29′E | NE | 16 | 4.1 | 66.0 | Andersson *et al.* (1998); Hyvönen *et al.* (2008) |

Table S2 ctnd.: Experimental sites used in this study. PFT abbreviations refer to BS = broadleaf deciduous, NE = needleleaf evergreen

| Site | Coordinates | PFT | Duration | Annual N | Total N | Reference |
|---|---|---|---|---|---|---|
| Lövnäs | 61°21′N/13°22′E | NE | 23 | 3.4 | 78.0 | Hyvönen *et al.* (2008) |
| Asa | 57°08′N/14°45′E | NE | 14 | 5.9 | 83.2 | Hyvönen *et al.* (2008) |
| Skogaby | 56°33′N/13°13′E | NE | 14 | 9.3 | 130.0 | Hyvönen *et al.* (2008) |
| Harplinge | 56°45′N/12°45′E | NE | 1 | 2.0 | 2.0 | Sikström (2002) |
| Össjö | 56°14′N/13°04′E | NE | 1 | 2 | 2 | Sikström (2002) |
| Norrliden N1 | 64°21′N/19°46′E | NE | 40 | 3.3 | 132.0 | Högberg *et al.* (2006) |
| Norrliden N2 | 64°21′N/19°46′E | NE | 40 | 6.6 | 264.0 | Högberg *et al.* (2006) |
| Norrliden N3 | 64°21′N/19°46′E | NE | 40 | 5.4 | 216.0 | Högberg *et al.* (2006) |
