## Supplementary material for "Whole-plant optimality predicts changes in leaf nitrogen under variable CO_2_ and nutrient availability": README

The file SI2_fertilisation_data_20190611.csv contains data from the literature on biomass and leaf N responses in N fertilisation experiments. For a full list of references, see SI1 Table S1 and S2.

Missing value: -999

Site: site name
PFT: plant functional type. NE = needleleaf evergreen, BS = broadleaf seasonal
Lon: longitude on a regular 0.5 x 0.5 degree grid for model runs. For the exact location of the site see SI1 Table S1 and S2
Lat:  latitude on a regular 0.5 x 0.5 degree grid for model runs. For the exact location of the site see SI1 Table S1 and S2
Harvest year: year in which the forest was set up as a plantation
Start year: year in which the experiment was started
N type: type of fertiliser added
Total N: Total amount of N added during the experiment, g N m-2
Annual N: Average amount of N added each year, g N m-2 yr-1
Duration: Duration of the experiment, years
Biomass: Biomass response to N addition, %
Biomass_std: Standard deviation of biomass response, where replicates were available, %
Biomass_year: Year when biomass measured
LeafN: Leaf N response to N addition, %
LeafN_std: Standard deviation of leaf N response, where replicates were available, %
LeafN_year: Year when leaf N measured
