## Supplementary material for "Whole-plant optimality predicts changes in leaf nitrogen under variable CO_2_ and nutrient availability": README~

The file SI2_fertilisation_data_20190611.csv contains data from the literature on biomass and leaf N responses in N fertilisation experiments. For a full list of references, see SI1 Table S1 and S2.

Site
PFT
Lon
Lat
Harvest
Start_year
N type
Total N
Annual N
Duration
Biomass
Biomass_std
Biomass_year
LeafN
LeafN_std
LeafN_year
